# Switchgrass Rhizosphere Metabolite Chemistry Driven by Nitrogen Availability

**DOI:** 10.1101/802926

**Authors:** Darian N. Smercina, Alan W. Bowsher, Sarah E. Evans, Maren L. Friesen, Elizabeth K. Eder, David W. Hoyt, Lisa K. Tiemann

## Abstract

Plants and soil microorganisms interact closely in the rhizosphere where plants may exchange carbon (C) for functional benefits from the microbial community. For example, the bioenergy crop, switchgrass (*Panicum virgatum*) is thought to exchange root-exuded C for nitrogen (N) fixed by diazotrophs (free-living N-fixers). However, this interaction is not well characterized and it is not known how or if switchgrass responds to diazotrophs or their activity. To explore this question, we assessed rhizosphere metabolite chemistry of switchgrass grown in a hydroponic system under two N levels and under inoculated or uninoculated conditions. We found switchgrass root exudate chemistry to be more responsive to N availability than to diazotroph presence. Total metabolite concentrations were generally greater under high N versus low N and unaffected by inoculation. Examination of rhizosphere chemical fingerprints indicates metabolite chemistry was also driven strongly by N availability with a greater relative abundance of carbohydrates under high N and greater relative abundance of organic acids under low N. We also found evidence of changes in rhizosphere chemical fingerprints by inoculation treatment suggesting a potential for switchgrass to respond or even recruit diazotrophs. However, we found little evidence of N treatment and inoculation interaction effects which suggests switchgrass response to diazotroph presence is not mediated by N availability.

## Introduction

Plants and soil microorganisms interact closely in the rhizosphere, where roots exude a variety of compounds including carbohydrates, organic acids, and amino acids, which influence the rhizosphere environment and the microbes that live there (Bertin et al 2003; Bais et al 2006; Jones et al 2009). Through these interactions, plants may exchange carbon (C) for functional benefits conferred by the soil microbial community, such as protection against pathogens or mobilization of soil nutrients (Bais et al. 2006; Jones et al. 2009). Root exudation is thought to be mainly driven by passive diffusion (Jones et al. 2009), yet more active routes such as anion channels and ATP-driven transport highlight the potential for plants to regulate both the amount and composition of exudates released (Bertin et al. 2003; Sasse et al. 2018). Through “modulation of their exudate profiles” (i.e. changes in chemical composition and abundance), plants can alter their rhizobiome (Sasse et al. 2018). Yet, it is still unclear if plants alter root exudate chemistry in response to the rhizosphere community’s functional capacity, such as the ability of the rhizobiome to fix nitrogen (N). We explored this question using the interaction between switchgrass (*Panicum virgatum*) and the known diazotroph (free-living N-fixing organism), *Azotobacter vinelandii* DJ, hereafter AV.

Switchgrass, a perennial C4 grass native to North America, is a promising cellulosic bioenergy crop favored for its high biomass production and climate change mitigation potential (Robertson et al. 2017). Switchgrass productivity is often unresponsive to fertilizer N additions, resulting in similar biomass yields with fertilizer N additions well above or below plant N demands (Parrish and Fike 2005; Ruan et al. 2016). Further, yields remain consistently high despite N removal via yearly harvest (Parrish and Fike 2005; Ruan et al. 2016). An N mass balance suggests that switchgrass is accessing an unaccounted-for N source at rates of 35 – 58 kg N ha^-1^ yr^-1^ (Roley et al. 2018). Switchgrass is also known to grow productively on low fertility marginal lands, which are often low in N (Robertson et al. 2017). This raises the question of how switchgrass is meeting its N demands.

One potential source of N for switchgrass is free-living N-fixation (FLNF) carried out by diazotrophs in the rhizosphere. There is strong evidence for this as FLNF has been measured in association with the switchgrass rhizosphere (Roley et al. 2018; Roley et al. 2019; Smercina et al. in press). In fact, Roley et al. (2018) found that FLNF could account for 80 – 100% of the estimated “missing” N. Switchgrass is also known to host a diverse community of diazotrophs in its rhizosphere (Bahulikar et al. 2014). Switchgrass-diazotroph interactions present a unique system for exploring plant response to rhizobiome functional capacity given their known association and high potential for switchgrass to benefit from FLNF carried out by diazotrophs.

In this study, we explored the interaction between switchgrass and AV under high and low N conditions. High N should be less favorable than low N for FLNF as increasing N availability is shown to generally downregulate FLNF (Reed et al. 2011; Smercina et al. 2019). We predicted that there would be observable rhizosphere metabolite changes in response to N availability and diazotroph presence. We first hypothesized that switchgrass root exudation would be greater under low N than high N conditions as switchgrass attempts to recruit and support diazotrophs and that this response will be amplified by the presence of diazotrophs. Second, we hypothesized that root exudates under low N, particularly when inoculated, would be dominated by C compounds selective of diazotrophs, such as organic acids (Baldani et al. 2014; Smercina et al. 2019). Lastly, we hypothesized that differences in root exudate chemistry would be driven by diazotroph presence over N availability.

## Materials and Methods

### Experimental setup

Switchgrass root exudates were collected from sterile switchgrass seedlings (var. Cave-in-Rock) grown hydroponically in sterile test tubes. To sterilize seeds, ∼250 seeds were placed in three, loosely capped 5 ml tubes inside a desiccator. In a fume hood, seeds were exposed to chlorine gas for 5.5 hours. Chlorine gas was generated by adding 3 ml of concentrated hydrochloric acid to 100 ml of 8.25% sodium hypochlorite. Seedlings were pre-germinated from sterile seeds on sterile filter paper in Petri dishes moistened with sterile Milli-Q water (Millipore, Burlington, MA, USA). Petri dishes of sterile seeds were incubated in the dark at 30°C for 5 days. Test tubes were prepared for planting with four Whatman #114 cellulose filter papers (GE Healthcare Life Sciences, Chicago, IL, USA) rolled into a ring inside the test tube such that roots would always grow between two layers of filter paper. Filter paper and test tubes were washed 3 times with HPLC-grade methanol prior to planting to remove potential contaminants and then autoclaved. Working inside a sterile biosafety cabinet with flame-sterilized forceps, one sterile switchgrass seedling was transplanted between two layers of filter paper in each test tube. After transplanting, 15 ml of a ¼ strength Hoagland’s nutrient solution (1.25 mM KCl, 1.25 mM CaCl_2_, 0.25 mM KH_2_PO_4_, 0.5 mM MgSO_4_, 0.012 mM H_3_BO_3_, 0.002 mM MnCl_2_*4H_2_O, 0.051 µM CuSO_4_*5H_2_O, 0.191 µM ZnSO_4_*7H_2_O, 0.124 µM Na_2_MoO_4_*2H_2_O, 2.7 µM NaFeEDTA) with either high or low N as NH_4_NO_3_ (described below) was added to the bottom of each tube. This facilitated wicking of nutrient solution up from the bottom, allowing roots to grow freely without being submerged in liquid. This also allowed diffusion of root exudates into the nutrient solution for later collection. After planting, each tube was immediately sealed with a gas permeable membrane (Breathe-EASIER membrane; Diversified Biotech, Dedham, MA, USA) to allow gas exchange while maintaining a sterile environment.

Nitrogen was added as ammonium nitrate at either a high N concentration (3.75 mM N-NH_4_NO_3_) or low N concentration (0.25 mM N-NH_4_NO_3_) with a total of 18 tubes per N treatment. These N concentrations were chosen based on preliminary experiments which demonstrated switchgrass N-deficiency at the low N and adequate N availability at the high N level (Friesen, unpublished data). An additional 18 tubes of each N treatment were prepared as described without planting to be used as background controls. All tubes were placed in a growth chamber on 16-hour day cycles (155 µmol m^-2^ s^-1^ light intensity) maintained at 60% humidity with 23°C daytime and 21°C nighttime temperatures. After 3 weeks of growth, half of all tubes (both N treatments and with/without plants) were inoculated with 1 ml of 10^7^ CFU ml^-1^ AV suspended in ¼ Hoagland’s solution described above. AV inoculum was cultured in LB broth at 24 °C for ∼16 hours until optical density (OD600) reached 0.1, corresponding to 10^7^ CFU ml^-1^ in exponential phase. AV cultures were then centrifuged at 3220 * *g* for 5 minutes, supernatant was discarded, and then cultures were resuspended in the same volume of sterile ¼ Hoagland’s solution prior to plant inoculation. Uninoculated seedlings received 1 ml of sterile ¼ Hoagland’s solution in place of inoculation. Plants were then grown an additional 3 days, with and without AV. This resulted in 4 treatment groups (High N, High N + AV, Low N, and Low N + AV) with and without plants for a total of 72 tubes (9 tubes per treatment). Sterility of all tubes was confirmed prior to inoculation by plating nutrient solution from each sample onto TY agar plates (5 µl each). Plates were placed in a 30 °C incubator for 3 days and then examined for growth.

### Sample harvest

Plant biomass and metabolite samples were harvested three days after inoculation. All harvest work was done in a sterile laminar flow hood with flame sterilized equipment. The porous membrane of each tube was removed and then the filter paper containing a seedling was carefully removed from the tube. Shoot biomass was clipped and root systems were carefully excised from the filter paper. Shoot and root biomass were dried at 60°C for at least three days to determine above- and belowground biomass, respectively. All nutrient solution from the bottom of the tube was transferred to a 50 ml conical tube. The filter paper was placed on 3 sterile pipette tips inside the same 50 ml tube and then the tube was centrifuged at 1610 * *g* for 5 minutes to collect residual solution from the filter paper. The solution was then transferred to a 15 ml conical tube and centrifuged at 3220 * *g* to pellet debris. Finally, the supernatant from 3 replicates per treatment group was collected and pooled for analysis, resulting in a total of 24 samples for NMR analysis, including 12 background control samples. Due to microbial contamination in one tube, only 2 samples were pooled for low N seedling treatment replicate 3. Samples were frozen at −80 °C until lyophilization.

### NMR analysis

Lyophilized material from each tube was weighed and resuspended in 180 µl of 0.5 mM dDSS (4,4-dimethyl-4-silapentane-1-sulfonic acid) in 10% D_2_O/ 90% H_2_O for NMR analysis. NMR spectra for all samples were collected using a Varian Direct Drive 600-MHz NMR spectrometer with a 5-mm triple resonance cold probe. NMR spectra were collected using standard Chenomx data guidelines (Weljie et al., 2006) employing a 1D ^1^H NOESY pre-saturation experiment and were processed, assigned, and analyzed using Chenomx NMR Suite 8.3 (Edmonton, AB, Canada). Spectra were quantified based on intensity relative to the internal standard. Sample metabolites were determined by matching chemical shift, J-coupling, and intensity against NMR signals of metabolites in the Chenomx library. Raw metabolite concentrations were reported as µM concentrations. It is important to note that not all compounds within the lyophilized material are detected via NMR and that a lack of detection does not mean an absence of the compound, but merely that the concentrations are below the NMR detection limit. Further, in this study we are specifically focused on C containing compounds.

### Data Analysis and Statistics

Raw metabolite concentrations were converted from µM to nmol g^-1^ dry material by first multiplying by the volume of dDSS used in resuspension (i.e. 1.8 x 10^-4^ L) to determine µmol of metabolite, then converting to nmol by multiplying this by 1000, and finally dividing by grams of dry, lyophilized material. Final compound concentrations, as nmol of detected metabolite g^-1^ dry material, were calculated by subtracting average compound concentration in background controls from samples under each N and inoculation treatment. Relative abundance was calculated as the nmol of detected metabolite g^-1^ dry material divided by total nmol of all detected metabolites g^-1^ dry material, thus relative abundance is nmol of detected metabolite nmol^-1^ total metabolites.

The distributions of metabolite concentrations were non-normal, requiring non-parametric statistical procedures. Therefore, comparisons of concentration and relative abundance of specific compounds by N treatment or inoculation were carried out using Friedman’s test, non-parametric two-way ANOVA in SAS (SAS v. 9.1). Principal components analysis (PCoA) was conducted in R using the *vegan* package tools *vegdist* and *adonis* (R version 3.6.0; Oksanen et al. 2019) with Bray-Curtis dissimilarity. Correlations of the dissimilarity matrix and compound group concentrations were carried out using the *vegan* tool *envfit.* We calculated 95% confidence ellipses for PCoA using *dataEllipse* in the *car* package (R version 3.6.0; Fox and Weisberg 2018).

## Results

### Metabolite concentrations

We successfully collected and analyzed the metabolite profiles of switchgrass and switchgrass with AV grown in a sterile, semi-hydroponic system. We detected and identified 34 rhizosphere metabolites representing eight compound groups including alcohols, amino acids/amino sugars, carbohydrates, cholines, ketones, nucleic acids, organic acids, and quinones (Table 1). Total measured rhizosphere metabolite concentration (nmol detected metabolite g^-1^ dry material) was on average 2x greater for high N plants than low N plants (F = 10.89, p = 0.0109; Fig. 1A). Though not significant, there was a trend towards greater detected metabolite concentrations with inoculation (F = 3.56, p = 0.0961; Fig. 1B). There was no significant interaction effect of N treatment and inoculation on total measured rhizosphere metabolite concentration.

**Figure 1.**
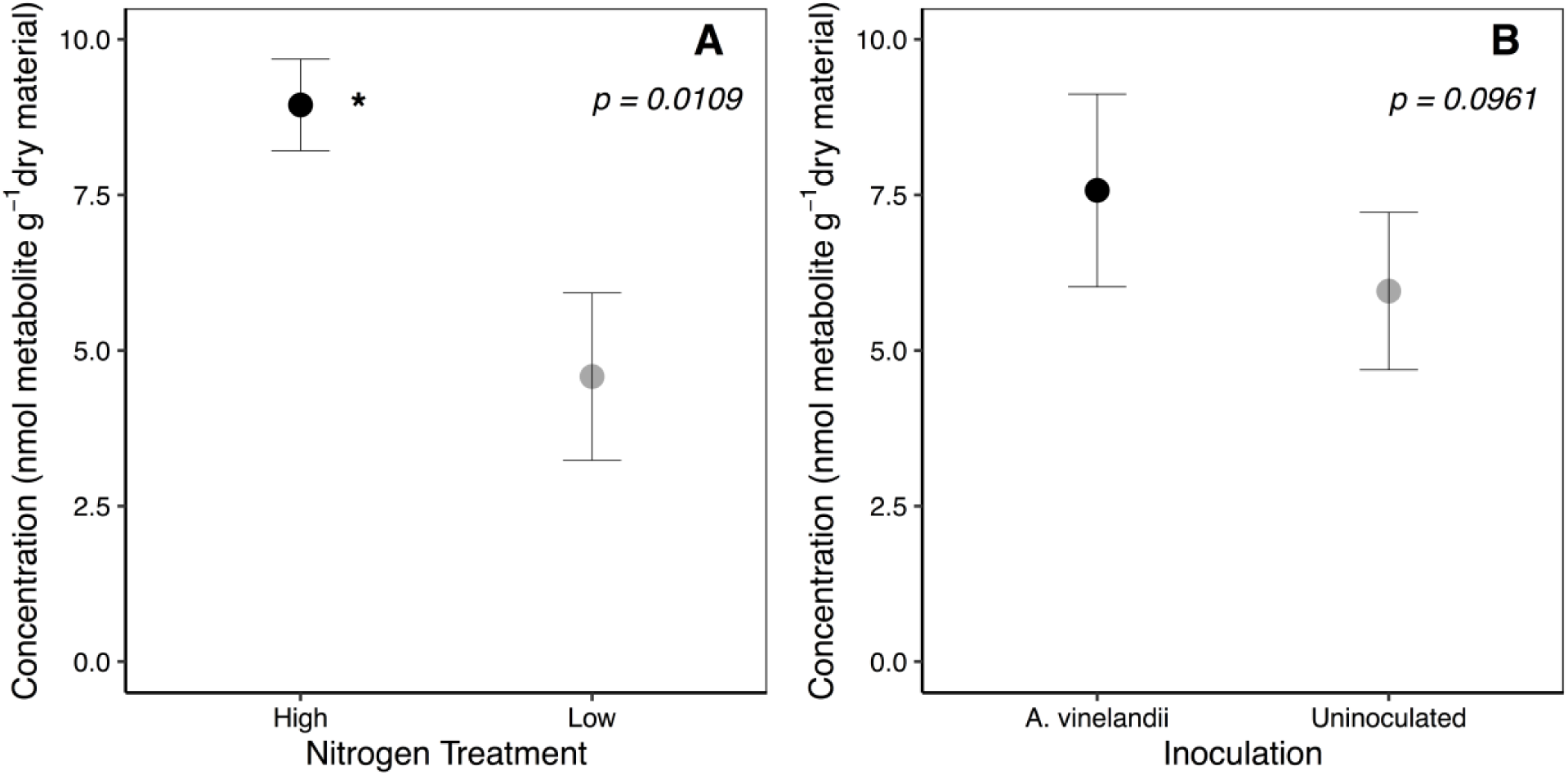
Concentration of detected total metabolite carbon by (A) nitrogen treatment and (B) inoculation treatment. Points represent average concentration +/− standard error (n = 6). Lower case letters indicate significant differences for p < 0.05. Differences in total metabolite carbon were not significant by inoculation treatment.

**Table 1.** Relative abundances (nmol detected metabolite nmol^-1^ total metabolites) of all compounds by treatment groups

| Compound Class | Compound | High N | High N + AV | Low N | Low N + AV |
| --- | --- | --- | --- | --- | --- |
| Alcohol | Ethylene glycol | 0.00804 ± 0.00413 | 0.00579 ± 0.00312 | 0.00691 ± 0.00536 | 0.00092 ± 0.00092 |
|  | **Methanol** <sub>N</sub> | 0.0026 ± 0.0017 | 0.0060 ± 0.0013 | 0.003 ± 0.003 | 0.0026 ± 0.0026 |
|  | Propylene glycol | 0.0418 ± 0.00875 | 0.0319 ± 0.00600 | 0.112 ± 0.0380 | 0.0665 ± 0.00579 |
| Amino Acid - Amino Sugar | 2 - Aminoisobutyric acid | 0.026 ± 0.011 | 0.01 ± 0.006 | 0.05 ± 0.05 | 0.03 ± 0.03 |
|  | Alanine | 0.014 ± 0.0047 | 0.016 ± 0.0036 | 0.03 ± 0.03 | 0.02 ± 0.009 |
|  | **Betaine** <sub>N<sup>†</sup></sub> | 0.00072 ± 0.00072 | 0.0677 ± 0.00365 | 0.0087 ± 0.0037 | 0.16 ± 0.056 |
|  | **Glycine** <sub>N</sub> | 0.040 ± 0.011 | 0.0086 ± 0.0057 | 0.028 ± 0.019 | 0.0016 ± 0.0011 |
|  | **Lysine** <sub>†</sub> | BDL | 0.14 ± 0.026 | BDL | 0.04 ± 0.04 |
|  | **Valine** <sub>N<sup>†</sup></sub> | BDL | 0.015 ± 0.00099 | 0.090 ± 0.041 | 0.002 ± 0.002 |
|  | Methylamine | 0.06 ± 0.06 | 0.04 ± 0.04 | 0.01 ± 0.01 | 0.01 ± 0.01 |
|  | Trimethylamine | BDL | BDL | BDL | 0.007 ± 0.004 |
| Carbohydrate | 1,3 - Dihydroxyacetone | 0.0080 ± 0.0051 | 0.003 ± 0.002 | 0.0015 ± 0.0015 | BDL |
|  | **Arabinose** <sub>N</sub> | 0.2031 ± 0.02590 | 0.171 ± 0.0524 | 0.108 ± 0.108 | 0.0285 ± 0.0167 |
|  | **Fructose** <sub>N</sub> | 0.219 ± 0.0257 | 0.218 ± 0.0697 | 0.07 ± 0.07 | 0.01 ± 0.01 |
|  | **Glucose** <sub>N</sub> | 0.1916 ± 0.01720 | 0.1001 ± 0.03810 | 0.0142 ± 0.0142 | BDL |
| Choline | Choline | BDL | 0.0013 ± 0.00069 | BDL | BDL |
|  | O-Phosphocholine | 0.01 ± 0.0022 | 0.011 ± 0.0042 | 0.021 ± 0.012 | 0.019 ± 0.0062 |
| Ketone | Acetone | 0.0297 ± 0.0163 | 0.0082 ± 0.0042 | 0.013 ± 0.013 | 0.034 ± 0.018 |
| Nucleic acid | Thymidine | 0.02 ± 0.009 | BDL | 0.004 ± 0.004 | 0.003 ± 0.003 |
|  | Thymine | 0.005 ± 0.005 | BDL | 0.01 ± 0.01 | BDL |
|  | **Uracil** <sub>†</sub> | BDL | 0.0387 ± 0.00810 | BDL | 0.270 ± 0.132 |
| Organic Acid | Acetate | BDL | BDL | 0.06 ± 0.03 | 0.02 ± 0.02 |
|  | Acetoacetate | 0.00875 ± 0.00444 | 0.0217 ± 0.00296 | 0.037 ± 0.023 | 0.014 ± 0.0098 |
|  | **Allantoin** <sup>N†</sup> | BDL | 0.010 ± 0.0051 | BDL | BDL |
|  | Benzoate | 0.01 ± 0.01 | BDL | BDL | BDL |
|  | Dimethylmalonic acid | 0.002 ± 0.001 | 0.0007 ± 0.0007 | 0.0092 ± 0.0055 | 0.0035 ± 0.0026 |
|  | Formate | 0.005 ± 0.005 | 0.006 ± 0.006 | 0.05 ± 0.05 | 0.02 ± 0.02 |
|  | Gallate | 0.025 ± 0.0051 | 0.019 ± 0.0033 | 0.076 ± 0.022 | 0.068 ± 0.032 |
|  | Glycerol | 0.022 ± 0.011 | 0.0246 ± 0.0126 | 0.06 ± 0.06 | 0.0814 ± 0.0432 |
|  | Isobutyrate | BDL | BDL | 0.01 ± 0.01 | 0.02 ± 0.02 |
|  | **Lactate** <sup>N</sup> | 0.01 ± 0.01 | 0.0269 ± 0.00280 | BDL | BDL |
|  | Succinate | BDL | BDL | 0.0298 ± 0.0298 | 0.00705 ± 0.00705 |
|  | Tartrate | 0.04 ± 0.02 | BDL | 0.09 ± 0.09 | 0.06 ± 0.06 |
| Quinone | Quinone | BDL | BDL | 0.003 ± 0.002 | 0.00092 ± 0.00092 |

BDL – Below Detection Limit. Compounds with significant treatment differences are bolded.
<sup>N</sup> Indicates significant N treatment effect
<sup>†</sup> Indicates significant Inoculum treatment effect
<sup>N†</sup> Indicates significant interaction effect

We explored differences in concentrations of specific compounds between N treatments and inoculation treatments (Table S1). Arabinose, fructose, glucose, glycine, lactate, and methanol were all significantly greater under high N and were not impacted by inoculation (Table S2). In contrast, lysine and uracil were significantly greater when plants were inoculated (p = 0.0043 and p < 0.0001, respectively) and were not impacted by N availability. Some compounds also had interactions between N treatment and inoculation, including allantoin, betaine and valine (Fig. 2; Table S2). Allantoin was greatest when plants were under high N and inoculated with no significant differences between high N, low N, or low N with inoculation (Fig. 2A, p < 0.0001). Betaine was also greatest in the high N with inoculation treatment followed by low N with inoculation, low N, and finally high N (Fig. 2B, p = 0.0123). Valine was greatest under low N without inoculation and high N with inoculation with no significant differences between high N and low N with inoculation (Fig. 2C, p < 0.0001).

**Figure 2.**
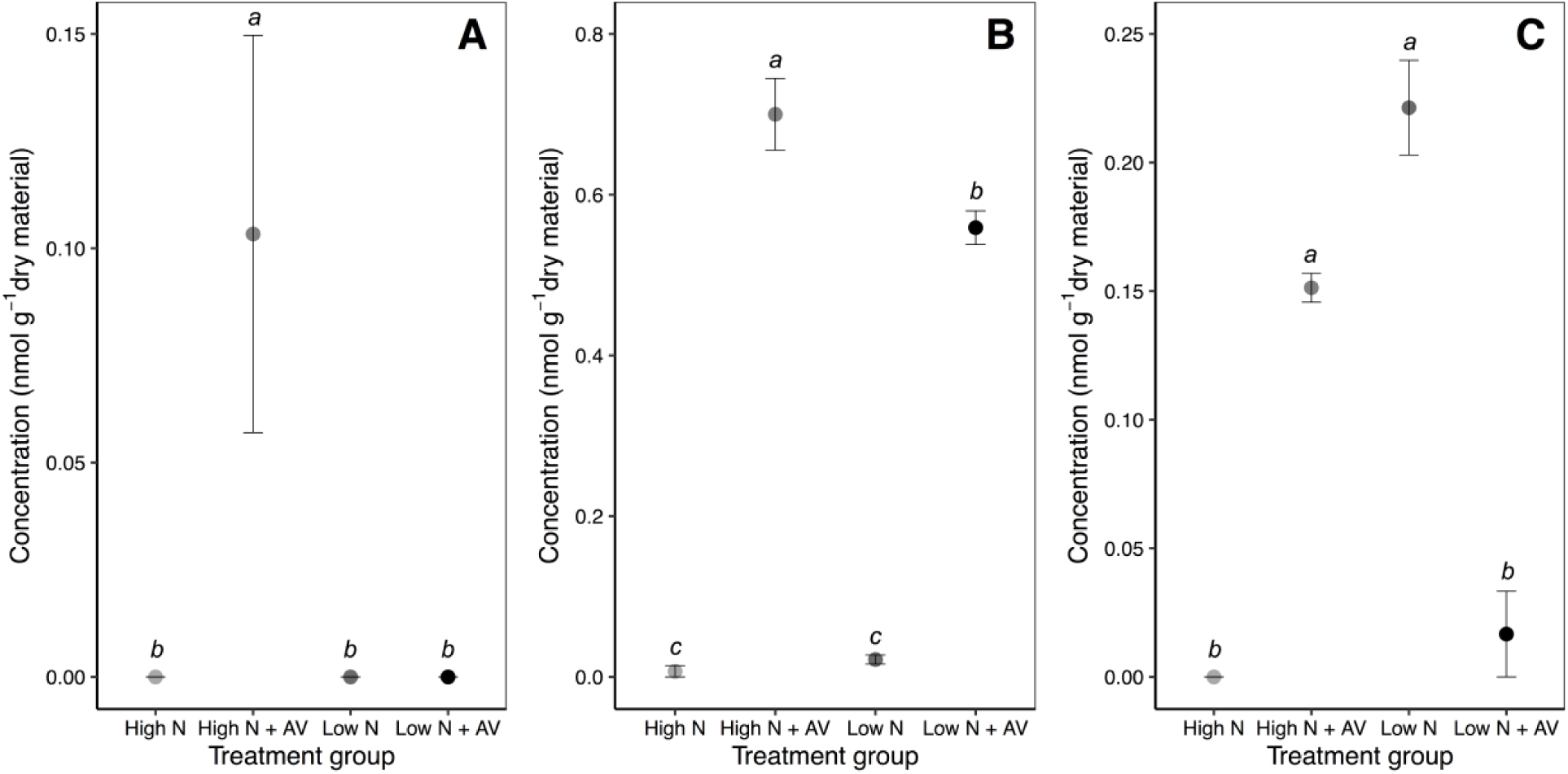
Concentration of (A) Allantoin, (B) Betaine, (C) Valine, compounds that had significant nitrogen by inoculation treatment effects. Points represent average concentration +/− standard error (n = 3). Lower case letters indicate significant differences for nitrogen by inoculation interactions where p < 0.05.

We also explored differences in concentration of compound classes across N and inoculation treatments (Table S3). Total concentration (nmol g^-1^ dried material) of alcohols and carbohydrates were significantly greater under high N (p = 0.0033 and p = 0.0009, respectively) while total choline concentrations were significantly greater under low N (p = 0.0055). Total amino acid and nucleic acid concentrations were greater when plants were inoculated with AV than when uninoculated (p = 0.0481 and p = 0.0038, respectively).

### Relative abundance

Because of differences in total metabolite concentration between N treatments, we also compared relative abundance of metabolite compound groups between N treatments (Fig. 3; Table S3). High N samples were dominated by carbohydrates with 55.1% of detected C compounds belonging to this compound group. Carbohydrates only represented 8.1% of detected C compounds in low N samples. We found low N samples to be dominated by organic acids with over 42.2% of detected C compounds represented by this group. Overall, high N samples had a greater abundance of carbohydrates (p = 0.0001) compared to low N samples. Low N samples had a greater abundance of cholines (p=0.0008) and organic acids (p = 0.0029) compared to high N samples. Carbon compounds detected in low N samples were more evenly distributed across compound groups with few differences between compound groups. Relative abundance of nucleic acids (p = 0.0144) was found to be significantly greater in inoculated rhizospheres. Similarly, amino acids relative abundance showed a marginal response (p = 0.0667), trending towards greater abundance with inoculation. Relative abundance of carbohydrates (p = 0.0382) was also significantly impacted by inoculation and was greater in uninoculated rhizospheres.

**Figure 3.**
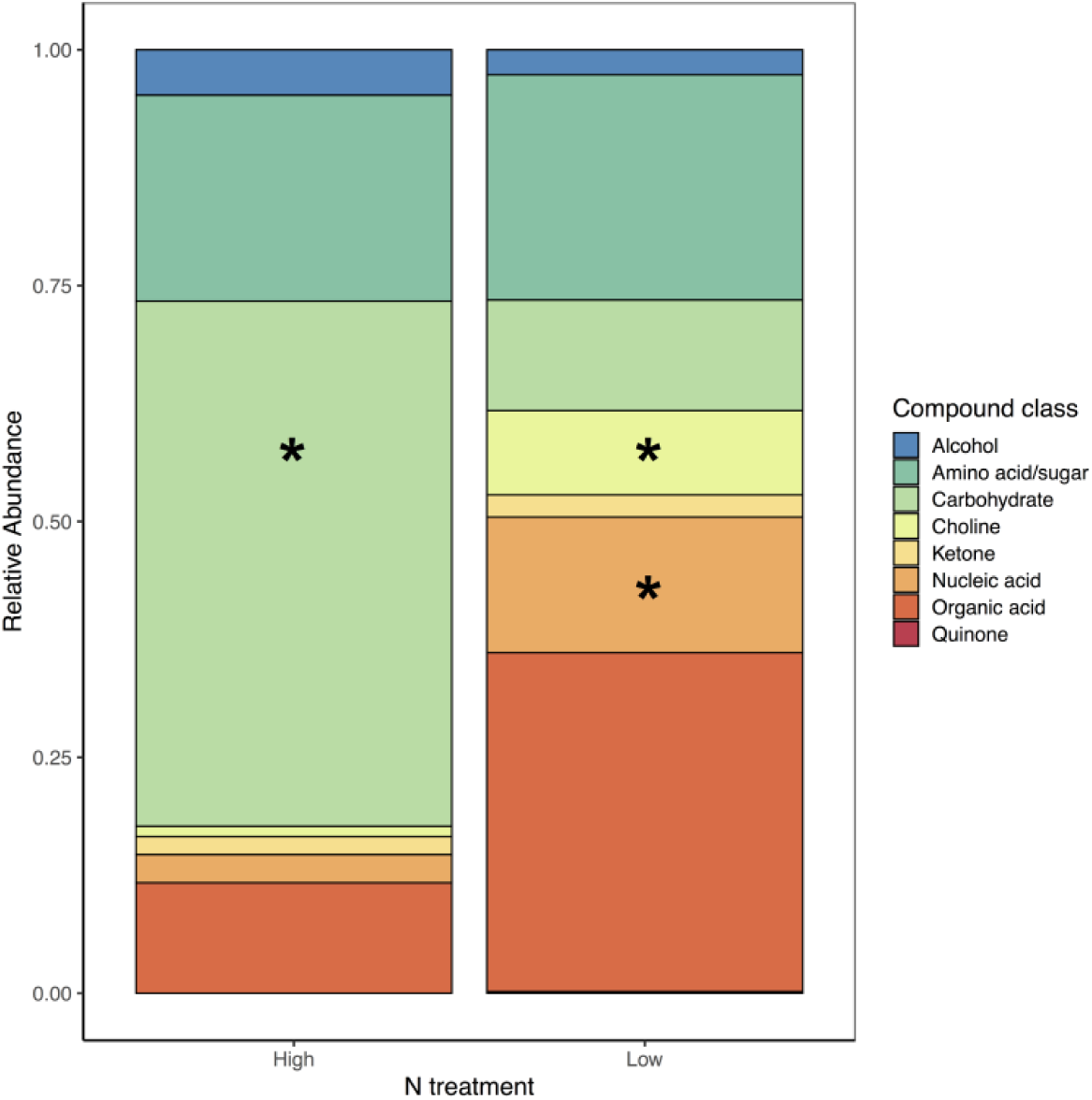
Relative abundance of compound groups by nitrogen treatment (n = 6 per nitrogen treatment). High N rhizospheres were carbohydrate dominated (55.1%), while low N rhizospheres were organic acid dominated (42.2%). Asterisk indicates significant difference between relative abundance at high versus low nitrogen where p < 0.05. Asterisks are placed on the bar with greater relative abundance.

### Rhizosphere metabolic fingerprint

We also examined the rhizosphere metabolite “fingerprints” based on relative abundance of all identified compounds in each sample (Table 1). Principal coordinates analysis (PCoA) shows fingerprints were driven by N treatment (R^2^ = 0.31096, p = 0.0004, Fig. 4) and inoculation (R^2^ = 0.16805, p = 0.0199), with no significant interaction. Environmental factor analysis indicates that spatial distribution in the PCoA is significantly correlated with carbohydrate (R^2^ = 0.7142; p = 0.005) and choline (R^2^ = 0.5931; p = 0.029) concentrations and marginally correlated with organic acid concentrations (R^2^ = 0.4854; p = 0.055).

**Figure 4.**
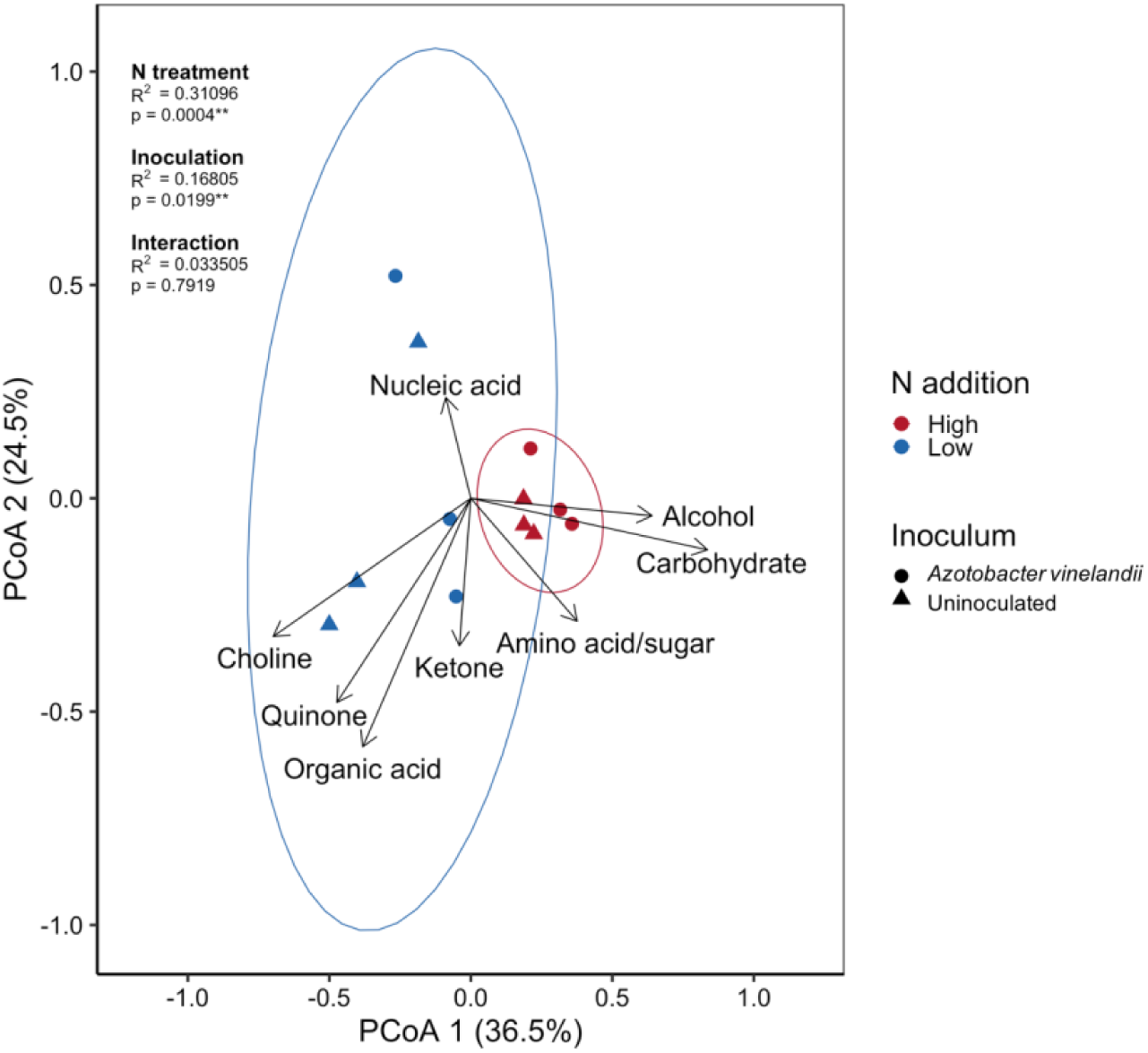
Principal coordinates analysis displaying Bray-Curtis dissimilarity based on relative abundance for all detected compounds. Ellipses represent 95% confidence ellipses by nitrogen treatments. 95% confidence ellipses for inoculation are not shown. Each point is represented by one sample. Overlaid vectors display correlations of the dissimilarity matrix with concentrations of each detected compound class.

### Plant Biomass

We found differences in shoot biomass by N treatment (p = 0.00503), but no differences in root biomass by N treatment (p = 0.259; Fig. 5). Shoot biomass was over 2.5x greater on average under high N than under low N. There were no differences in shoot or root biomass by inoculation treatment.

**Figure 5.**
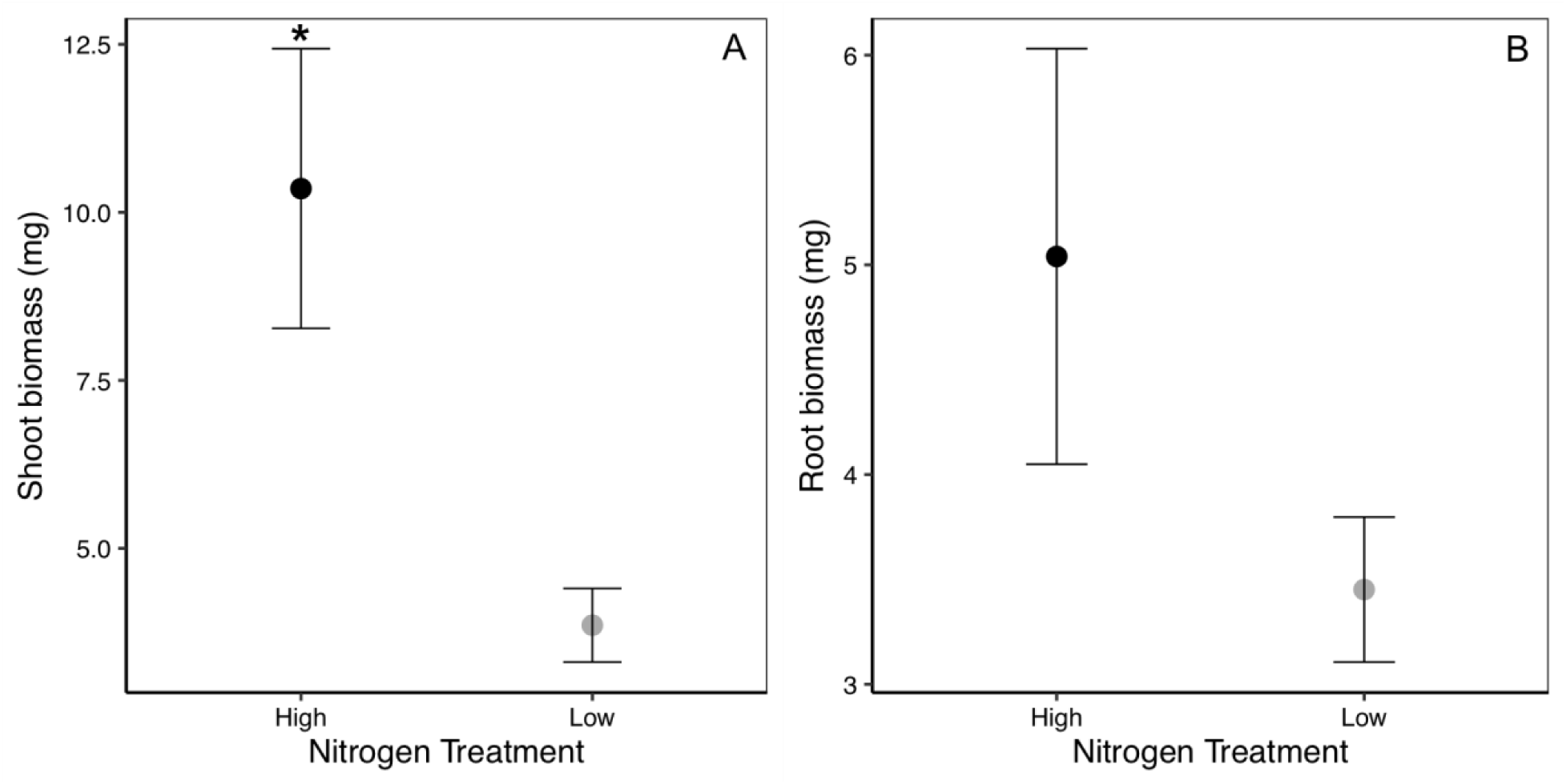
Shoot (A) and root (B) biomass by nitrogen treatment. Shoot and root biomass are not presented by inoculation treatment because results were not significant. Points represent average biomass +/− standard error (n = 6). Asterisk indicates significantly greater biomass by nitrogen treatment. Root biomass was not significantly different between nitrogen treatments.

## Discussion

Our results indicate that N availability exerted more control over rhizosphere metabolites than diazotroph presence. N availability was the main driver of total root exudation with high N plants exuding 2-fold greater total detected C compounds. We also found high N availability resulted in 2.5x greater shoot biomass while root biomass was not significantly different between N treatments, suggesting that increased exudation may be the result of greater photosynthetic activity and aboveground productivity rather than increased root biomass. We found no differences in shoot or root biomass with inoculation treatment, however this may indicate that roots were not exposed to inoculation long enough to see a large response in growth. Total metabolite concentration also did not show a response to inoculation. This was surprising as we hypothesized that switchgrass would increase exudation under low N in order to recruit and/or support diazotrophs. We also hypothesized that inoculation would further increase the differences in exudation by N treatment, but found no such result for total exuded C compounds. It is important to note that plants in our study had a short exposure to diazotrophs (i.e. 3 days) which may have limited any detectable responses in C quantity to diazotroph presence. Further, we were unable to assess inoculum survival and therefore, there is potential for reduced survival of the inoculum which may have also limited detectable responses to diazotroph presence. However, total quantity of exuded C may not be as important to diazotroph activity as C chemistry (Smercina et al. 2019) and indeed in this study, we find that changes in metabolite chemistry provide more insight into the switchgrass-diazotroph interaction than total exuded C.

Consistent with our second hypothesis we found that low N rhizospheres had a greater relative abundance of organic acids compared to high N rhizospheres. Organic acids are the preferred C source of many diazotrophs and are often used to isolate diazotrophs from soils (Baldani et al. 2014; Mukhtar et al 2018; Smercina et al. 2019). The greater relative abundance of organic acids observed under low N could indicate the ability of switchgrass to support diazotrophs through increased exudation of preferable C sources. However, we found no significant differences in organic acid concentration or relative abundance with inoculation or N*inoculation effects indicating that any changes in organic acid exudation by switchgrass may not be directly in response to diazotroph presence. We also found high N rhizospheres were dominated by carbohydrates with over 55% of detected metabolites coming from this compound group, while carbohydrates represented just over 8% of metabolites detected in low N rhizospheres. We did not find any significant differences in carbohydrate concentration by inoculation treatment or N*inoculation effects, but found uninoculated rhizospheres to have a greater relative abundance of carbohydrates than inoculated rhizospheres. These observed differences in carbohydrate concentrations by N treatment most likely resulted from greater photosynthetic activity in the high N plants which had greater aboveground biomass rather than a response to diazotroph presence. Interestingly, the differences in carbohydrate relative abundance by inoculation treatment may indicate that AV was actively consuming exuded carbohydrates. Although there is evidence that organic acids are a preferable C source for some diazotrophs, including AV (Mukhatar et al. 2018), we did not find evidence of organic acid uptake by AV (i.e. no significant difference in organic acid relative abundance by inoculation treatment).

Using the relative abundance of all compounds, we generated rhizosphere metabolite profiles, or “fingerprints.” Exploring differences in fingerprints is likely to provide the most information about plant-diazotroph interactions by allowing us to directly compare overall differences in metabolite chemistry across treatments. We found that fingerprints differ significantly between N treatments and inoculation treatments, but with no significant interaction between the two. A lack of fingerprint response to the interaction of N treatment and inoculation indicates that switchgrass response to diazotroph presence was not influenced by N availability, even though metabolite chemistry responded to changes in N and diazotroph presence independently. We also find that overall metabolite profiles are correlated with carbohydrate and choline concentrations and marginally with organic acid concentrations. Collectively, these results tell us that switchgrass may respond to reduced N availability by attempting to recruit diazotrophs (i.e. increased organic acid exudation). Though we find no strong evidence in this study that switchgrass is directly supporting diazotrophs, we do find evidence for potential pathways for direct switchgrass-diazotroph interaction.

Concentrations of most compounds in our study showed no significant response to inoculation treatment, thus limiting evidence for direct switchgrass-diazotroph interaction. However, we did find responses in both uracil and lysine concentrations, two compounds potentially important to plant-microbe interactions. Uracil could simply be of microbial origin; previous work shows that uracil is actively excreted from N-fixing bacteria (*Azotobacter agilis* B 170; Katayama et al., 1967), but it is also known to be important in the production of small, non-coding RNAs involved in plant-microbe interactions (Weiberg et al. 2014). Therefore, uracil may be indicative of direct switchgrass-diazotroph interaction. Similarly, lysine is a precursor to glutamate, an important signaling compound for plant growth and for plant response to environmental conditions, particularly stress responses like osmotic-stress (Arruda et al. 2000; Galili 2002). Lysine is also produced in large amounts by *A. vinelandii* strains (Gonzalez-Lopez et al. 1983) and has been shown to be essential for ATP binding to the iron co-factor of nitrogenase synthesized by *A. vinelandii* (Seefeldt et al. 1992). It is plausible that the lysine detected here was produced by the AV inoculum and may indicate active N-fixation by AV. Further, this production of lysine by AV may have induced stress responses in switchgrass, which is further evidenced by detection of compounds like allantoin, betaine, and valine.

Allantoin, betaine, and valine have all been associated with plant stress response, particularly osmotic stress response. All three compounds were detected in this study and were found to have significant N treatment by inoculation responses. Allantoin concentration under high N + AV was over two-fold greater than the other three treatment combinations. Allantoin has been shown to play a role in osmotic stress response of *Arabidopsis* (Watanbe et al. 2013). It is plausible that allantoin production was induced because of lysine production by AV or because of the hydroponic growth system. Allantoin presence could also be indicative of phosphorus deficiency (Tawaraya et al. 2014). This is plausible in our system, particularly in the high N + AV treatment, as competition for nutrients between plants and microbes may result in P limitation relative to N causing stoichiometric imbalances (Sterner and Elser 2002). Allantoin may also be indicative of active N recycling from microbial biomass as it is also an intermediate product of purine catabolism (Watanbe et al. 2014). Greater biomass (both plant and potentially microbial) in the high N + AV treatment would result in greater availability of biomass-derived N and therefore, we could expect greater allantoin production through the purine catabolism pathway.

Betaine was also detected in the rhizosphere and may indicate plant and microbial stress response (Sakamoto and Murata 2002; Wargo 2013). In plants, betaine production can occur in response to high salt environments (Sakamoto and Murata et al. 2002), while in bacteria, betaine is an important osmoprotectant (Wood 2011, Borton et al. 2018). Betaine concentrations were generally greater with inoculation, but showed inverse effects of N treatment where betaine concentration was greater in high N when inoculated, but greater in low N when uninoculated. This suggests that betaine was produced by both plants and microbes in our system. While some microbes convert betaine to glycine which would provide a pathway for N cycling (Wargo 2013), it seems most likely that betaine production was a stress response as concentrations were generally greater where we might expect greater plant and microbial biomass.

Lastly, valine was detected in our system and showed complex treatment responses which may indicate multiple drivers of valine production. When uninoculated, valine production most likely indicates stress response in plants, potentially osmotic stress (Gargallo-Garriga et al. 2018). However, when inoculated, valine may also indicate biofilm formation by the inoculum (Valle et al. 2008). Our pattern of results for valine suggests a combination of the two where uninoculated plants in low N conditions exhibit strong stress response which is alleviated under high N, and inoculum are forming biofilms under high N conditions, but not as strongly under low N conditions. Collectively, the presence of allantoin, betaine, and valine suggest plants may have been responding as if osmotically stressed, though this response was altered by N availability and inoculation.

In conclusion, our findings indicate that switchgrass rhizosphere metabolite chemistry is more responsive to N availability than presence of a diazotroph inoculum. Plant responses to diazotroph presence may have been dampened by short exposure time and potentially reduced inoculum survival. Overall greater detected C compounds under high N was counter to our hypothesis that switchgrass would exude more C when N-limited and indicates that aboveground productivity rather than plant recruitment of diazotrophs is a strong driver of exudation. Metabolite chemistry was also driven more strongly by N availability with a greater relative abundance and concentration of carbohydrates under high N and greater relative abundance of organic acids under low N. These changes in chemistry suggest a potential for switchgrass to recruit diazotrophs when N limited, but that this response is likely driven by plant N demands over interactions between switchgrass and diazotrophs.

## Acknowledgements

This work was conducted under the MMPRNT project, funded by the DOE BER Office of Science award DE-SC0014108. A portion of research was performed under the Facilities Integrating Collaborations for User Science (FICUS) program and used resources at the DOE Joint Genome Institute and the Environmental Molecular Sciences Laboratory (grid.436923.9), which are DOE Office of Science user facilities. Both facilities are sponsored by the Office of Biological and Environmental Research and operated under contract numbers DE-AC02-05CH11231 (JGI) and DE-AC05-76RL01830 (EMSL). In particular, the metabolomics profiling measurements were made using EMSL instrumentation (Proposal ID: 49977).

